# Ready, set, yellow! Color Preference of Indian Free-ranging Dogs

**DOI:** 10.1101/2024.02.01.578151

**Authors:** Anamitra Roy, Aesha Lahiri, Srijaya Nandi, Aayush Manchalwar, S Siddharth, J V R Abishek, Indira Bulhan, Shouvanik Sengupta, Sandeep Kumar, Tushnim Chakravarty, Anindita Bhadra

## Abstract

Most of the research on color vision related behaviors in dogs has involved training the dogs to perform visual discrimination tasks. We investigated the meaning of color to untrained Indian free-ranging dogs (FRDs). Using one-time multi-option choice tests for color preference in 134 adult dogs, we found the dogs to prefer yellow objects over blue or gray ones (p < 0.001, Cramer’s V = 0.433) while there was no preference between blue and gray (N = 102, p = 0.165). We next pitted a yellow object against a gray object that had food. Here, the dogs ignored the food to approach the yellow object first, both when the food was biscuit (N = 52, p < 0.001, Cramer’s V = 0.576) and chicken (N = 61, p < 0.001, Cramer’s V = 0.540), indicating the color preference to be quite strong. Color preference has previously been investigated in many other animals and has implications for behaviors like mate choice and foraging. Our study provides a new perspective into the ecology of Indian FRDs and might have implications for pet dogs as well, if they too show this preference.

**Highlights:**

- Indian free-ranging dogs (FRDs) show preference toward the color yellow over blue and gray.
- Indian FRDs show no preference between blue and gray colors.
- Attraction towards a yellow object can be stronger than attraction towards food rewards for Indian FRDs.

## 1. Introduction

Color vision and color preference have been subjects of research interest for over a century (reviewed by Staples, 1931). Decades of research have confirmed color vision in most vertebrates and arthropods (Kelber, Vorobyev, & Osorio, 2003, Baden & Osorio, 2019, Yilmaz et al., 2022). The presence of color vision in a species does not automatically suggest the presence of color preference, but to have color preference, the faculty of color vision is essential. There are multiple biological underpinnings for color preference, which may vary from species to species, while color vision essentially requires just the presence of cone cells (Kelber et al., 2003). Color preference has been found in many animals across myriad taxa. Humphrey (1971 & 1972) investigated color preference of *Rhesus* monkeys and found the monkeys to demonstrate a preference for different hues and brightness of colored fields just like they prefer the content of one picture over another. Humphrey (1972) suggested interest (towards the content of stimulus pictures) and pleasure (reaction to hue and brightness) to be the two drivers of the monkeys’ visual preference. Preference towards colors on an animal’s body is often linked with mate choice (reviewed by Higham & Winters, 2016). Burley (1982, 1986) found zebra finches to prefer conspecific body colors (species confidence) in their leg bands, which affected the birds’ mate choice. However, Wang et al. (2018) noted that zebra finch color preference has had poor replication across studies, and hitherto observed significant results might not be generalizable at species level.

Ecological valence or the usual role of a particular color in an animal’s life, has been suggested as another driver for color preference (Palmer & Schloss, 2010). Several studies have explored the psychological impact of colors which are often linked with ecological valence (reviewed by Elliot, 2015). It is postulated that the redness of primate faces being an indicator of health and mood might have been a driver for primates developing red-green color vision (Changizi, Zhang, & Shimojo, 2006). Some color preferences like that of bowerbirds seem to be culturally transmitted and thus vary across populations of the same species (Diamond, 1986).

In dogs (*Canis lupus familiaris*), color vision was established by Neitz et al. (1989) who characterized their cone cell activities to peak at 429nm and 555nm, indicating a dichromatic color vision (blue-yellow spectrum) similar to red-green colorblind humans. Further behavioral study (Siniscalchi et al. 2017), electroretinogram photometry (Jacobs et al. 1993), and topographical study of retinal cone cells (Mowat et al. 2008) have solidified our understanding of dog color vision and it is possible to digitally simulate a dog’s vision (Pongrácz et al., 2017: https://dog-vision.andraspeter.com/). However, all behavioral research related to canine color cognition so far has been focused on color discrimination tasks after a period of training: to mainly understand the limits of canine color vision (Tanaka et al. 2000, Byosiere 2018, Kasparson et al. 2013). It was hitherto unknown if and how untrained dogs are affected by color cues in their environment. The only available research on this topic is a non-peer-reviewed article on canine color preference regarding chew toys by Wong (2007) where the dogs had prior experience with chew toys of unreported colors.

The question of usefulness of color cues is especially important for FRDs, as unlike pets, they are not under direct human supervision (Serpell, 1995), and they need to make numerous important decisions daily in order to survive. It is possible that each available cognitive ability, including color vision, is important to these dogs even though their sense of smell is highly sensitive (Beaver, 2009; Hayes et al., 2018; Kokocińska-Kusiak et al., 2021). Notably, olfactory cues are limited by wind directions and the slow nature of diffusion (Fick, 1885). Visual cues thus might be much more reliable for a quick introductory assessment of an object. Indian FRDs are primarily scavengers of human-generated waste (Vanak and Gompper, 2009) from dumps that are often overwhelmingly smelly (personal observation) and can demand utilization of visual cues. There already is experimental evidence (Polgar et al., 2015) that pet dogs rely on different senses at different distances from an object of interest. Moreover, previous work on FRDs suggests that they use both visual and olfactory cues during scavenging, though the decision to eat something is mostly arrived at using a Rule of Thumb “If it smells like meat, eat it” (Bhadra et al., 2015, Sarkar et al., 2019).

We carried out choice tests on FRDs in India to understand if they have any color preferences. Further, to ascertain whether this preference affects the dogs in their usual lives at all, we tested the strength of their color preference by pitting their preferred color against a food reward. If dogs ignore food and approach their preferred color instead, they risk losing a vital resource.

## 2. Materials and methods

### 2.1. Subjects, locations, and time

The experiments were conducted in urban, semi-urban, and rural locations in Nadia and North 24 Parganas districts, and the city of Kolkata in West Bengal, India (details available in Online Resource 8). A total of 458 visibly fit adult FRDs were included in the experiments. The sexes of the dogs were noted visually according to their genitals. To avoid learning bias due to repetitions, each dog was tested only once. We chose new experiment locations each day to avoid repeating dogs by mistake. All trials were done in daylight and all the options (bowls) in one trial were placed equally under direct sunlight, or equally in shade to ensure they were under the same lighting conditions.

### 2.2. Experimental setup

#### 2.2.1. Three-choice experiment (yellow, blue, gray)

The experimental setup (Fig. 1) consisted of three terracotta bowls of similar size (diameter 12.58 ± 0.46cm, height 4.70 ± 0.23cm) and shape, each colored differently by us using Fevicryl^®^ Acrylic paint. Three colors were used: golden yellow, Prussian blue, and gray (manually created by mixing black and white). These bowls were then placed on a 70cm x 17cm cardboard platform made waterproof with brown packing tape.

**Fig. 1.**
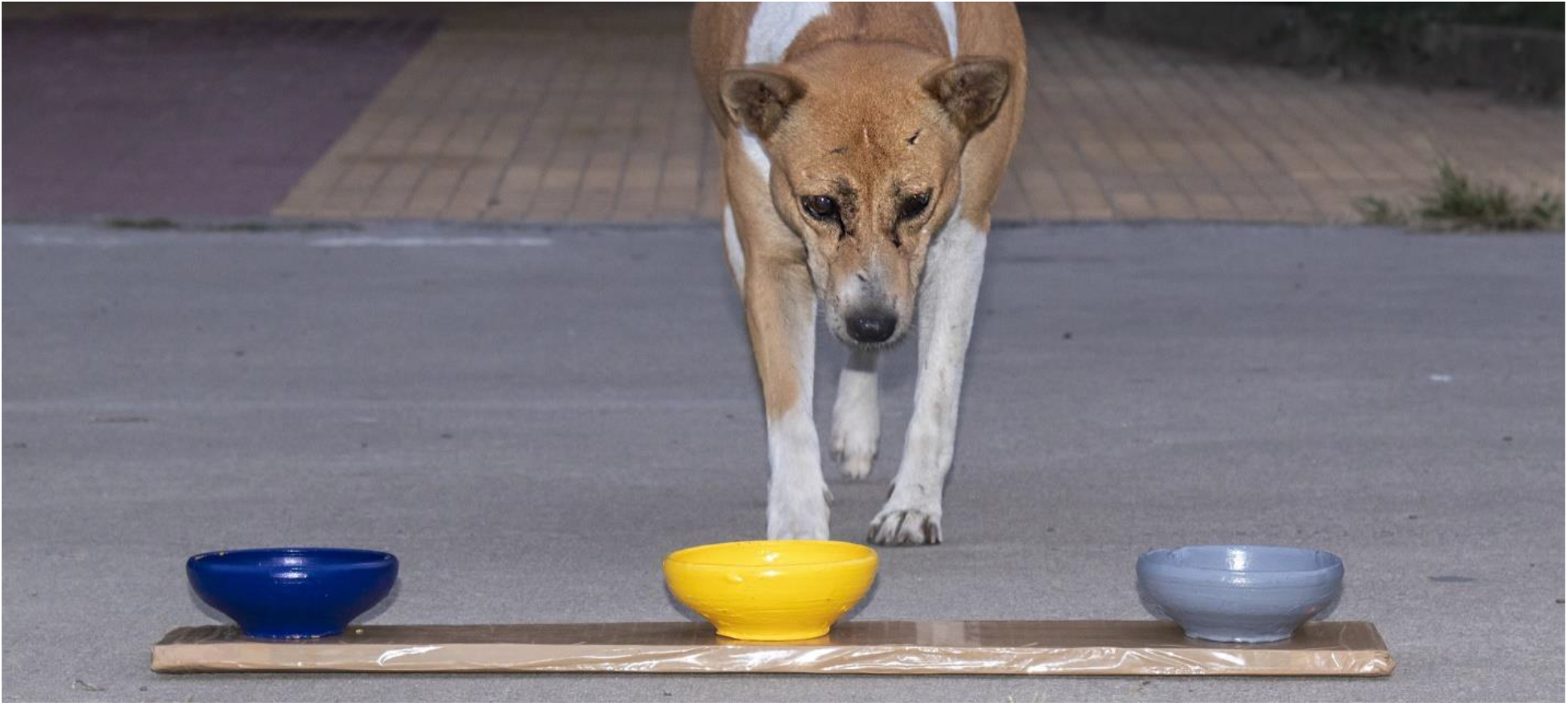
Experimental setup for the three-choice experiment. Photo: Sourabh Biswas.

In the “with_food” condition, each bowl had an equal amount of food in the form of a 1/4^th^ piece of a Brittania^®^ Marie Gold^®^ biscuit. In the “no_food” condition, no bowl had any food. See Online Resource 1 for video of the experiment.

#### 2.2.2. Two-choice experiment (blue, gray)

The three-choice experiment showed yellow to be the most preferred color (see results, section 3.1). Next, to identify any second preference, a two-choice test was done between blue and gray. The same blue and gray bowls as described earlier were used. We did not use the cardboard platform here since experimenter 1 could hold each bowl in their hands. All the trials were of “with_food” nature. See Online Resource 2 for video of the experiment.

#### 2.2.3. Control for the smell of paint

Since different paints (although the same brand and thus hopefully the same volatiles) were used to paint the bowls, it was important to make sure that the dogs were making their choice based on the visual cue of the paint and not the olfactory cue. Therefore, we painted two sets (blue and yellow) of terracotta bowls only on the bottom and covered them with plastic sieves (Fig. 2). This way we made sure all visual cues of the paint were blocked, but the smell could still permeate through the porous sieve. The covered bowls were placed on the same cardboard platform. See Online Resource 3 for video of the experiment.

**Fig. 2.**
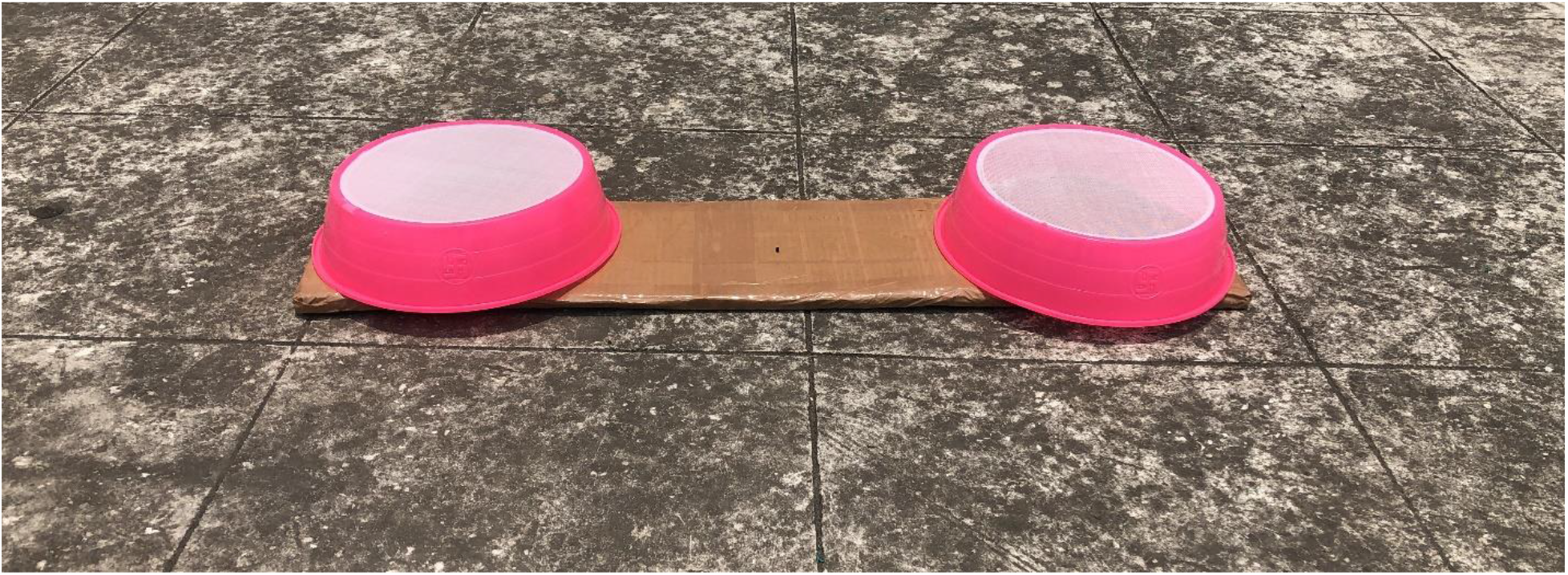
Experimental setup for the control for the smell of paint experiment

#### 2.2.4. Strength of preference experiment (“with_food” vs “no_food”)

The strength of preference experiment was devised to compare the dogs’ attraction towards yellow, against their attraction towards food which was associated with gray, a non-preferred color. The same bowls as before were used for this experiment, but to ensure the highest visibility of food, the bowls were upturned, and the food was placed on top of the upturned bowl.

##### 2.2.4.1. Control

The piece of biscuit being used in our experiments was quite small (a quarter of a circle with approximately 3 cm radius) and while FRDs do eat carbohydrates, they prefer meat (Bhadra et al., 2015, Sarkar et al, 2019). Thus, for the trials to work as intended, it was important to ensure that the dogs indeed prefer to choose the biscuit over an empty bowl when the color of the two bowls are the same. We chose both bowls to be yellow because the brownish biscuit is most camouflaged against the yellow bowl and thus if the dogs can identify and choose it, they should have no problem choosing it when it is atop a more contrasting gray bowl. Thus, the two options for the control trials were both yellow bowls, only one of which had a piece of biscuit on it.

##### 2.2.4.2. Trials

In the trials, we kept a piece of food on the gray bowl in a similar fashion to 2.2.4.1, while the yellow bowl was kept empty, thereby offering the dogs a choice between food and their preferred color. There were two conditions: a) where the food was a piece of biscuit as mentioned before, and b) where the food was a ∼15g piece of raw chicken. See Online Resource 4 and 5 for videos of the experiment.

### 2.3. Experimental protocol

The general protocol followed the one-time multi-option choice test (OTMCT) module described by Bhadra & Bhadra (2014). While traveling on the streets of the aforementioned locations, when an FRD was located, experimenter 1 approached the dog with the experimental setup. At about 1.5-2 meters from the dog, the experimenter vocalized positively towards the dog and placed the setup on the ground in a way that would keep the options equidistant to the dog. Then, experimenter 1 slowly backed away 5-10ft and assumed a neutral pose. Experimenter 2 videographed the whole experiment for analysis at a later time. A trial was deemed unsuccessful if the dog did not initiate an approach within 1 minute, or if the dog showed clear disinterest (e.g., going away from the area, closing its eyes and lazing). In case multiple dogs were present nearby, a trial was considered successful if no other dog came near the focal dog or the setup before the choice was made.

In each trial, the order of the bowls was randomized, and all trials happened with fresh bowls, i.e., the bowls were cleaned after each trial to remove any scent from dogs before being used again in a trial. Whichever bowl a dog investigated (nose within about 5cm of a bowl) first, was taken as that dog’s choice.

## 3. Results

All statistical analyses were performed with R (R core team, 2022) in RStudio (RStudio, 2019). The following packages were used for analysis and presentation: ggplot2 (Wickham, 2016), data.table (Barrett, Dowle, & Srinivasan, 2023), and DescTools (Signorell, 2023).

### 3.1. Three-choice experiment (yellow, blue, gray)

We successfully tested 76 dogs (50 males, 26 females) for the “with_food” and 58 dogs (27 males and 31 females) for the “no_food” condition. No difference was observed for the first choice between these conditions (Table 1; contingency *χ*^*2*^= 0.219, N= 134, df = 2, p = 0.896), and thus, the data were pooled for further analysis. Overall, yellow emerged as the most preferred color (Fig. 3, goodness of fit *χ*^*2*^ = 25.134, N = 134, df = 2, p < 0.001, Cramer’s V = 0.433).

**Fig. 3.**
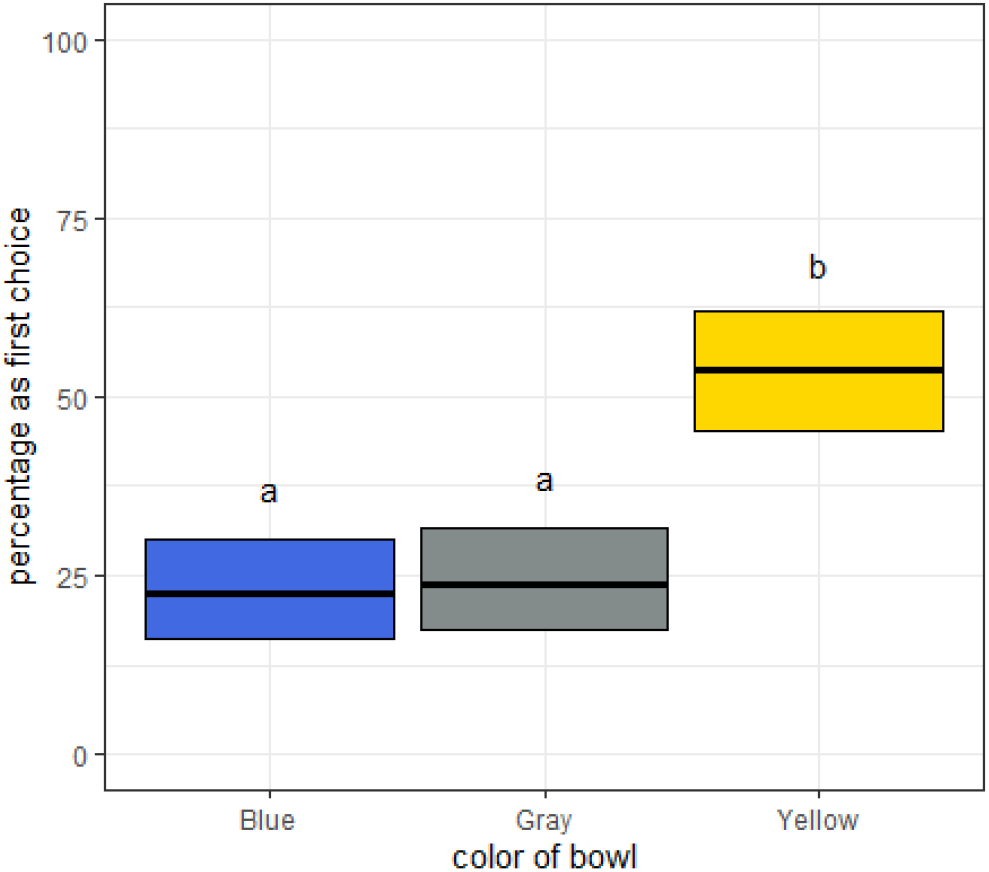
A box plot showing the percentage of times each color (Blue, Gray, and Yellow) was chosen the first time by a dog in the three-choice experiment. The central bold lines in the boxes are the observations, around which the boxes represent Wilson’s 95% CI. Different letters atop boxes denote significant statistical differences.

The preferred color was not different for male and female dogs (Data: Online Resource 7, contingency *χ*^*2*^= 1.938, df = 2, p = 0.379). The relative positions of the bowls had no significant effect on the choice of color (Data: Online Resource 7, contingency *χ*^*2*^ = 3.884, df = 4, p = 0.4218).

### 3.2. Two-choice experiment (blue, gray)

We successfully tested 102 dogs (51 females, 46 males, and 5 dogs of unknown sex) where blue was chosen 44 times and gray was chosen 58 times. There is no significant preference between these colors (goodness of fit *χ*^*2*^ = 1.921, df = 1, p = 0.165).

### 3.3. Control for the smell of paint (covered yellow, covered blue)

We successfully tested 55 dogs (29 females, 26 males). Blue was chosen 32 times and yellow was chosen 23 times. The observations do not significantly differ from chance (goodness of fit *χ*^*2*^= 1.472, df = 1, p = 0.224).

#### 3.4.1. Control for strength of preference (yellow with food = biscuit, yellow with no food)

We successfully tested 54 dogs (20 females, 33 males, and 1 dog of unknown sex). The bowls with food (biscuits) were chosen 35 times while bowls with no food were chosen 19 times. The preference is significant (goodness of fit *χ*^*2*^ = 4.740, df = 1, p = 0.029, Cramer’s V = 0.296).

#### 3.4.2. Strength of preference (gray with food, yellow with no food)

We successfully tested 52 dogs (20 males, 32 females) with biscuits as food (Fig 4). The gray + food combination was chosen 11 times while the empty yellow bowl was chosen 41 times (goodness of fit *χ*^*2*^ = 17.308, df = 1, p < 0.001, Cramer’s V = 0.576). When the food was ∼15g of chicken, the empty yellow bowl was chosen 47 times out of 61 successful trials (28 males, 33 females) (goodness of fit *χ*^*2*^ = 17.852, df = 1, p < 0.001, Cramer’s V = 0.540).

**Fig 4.**
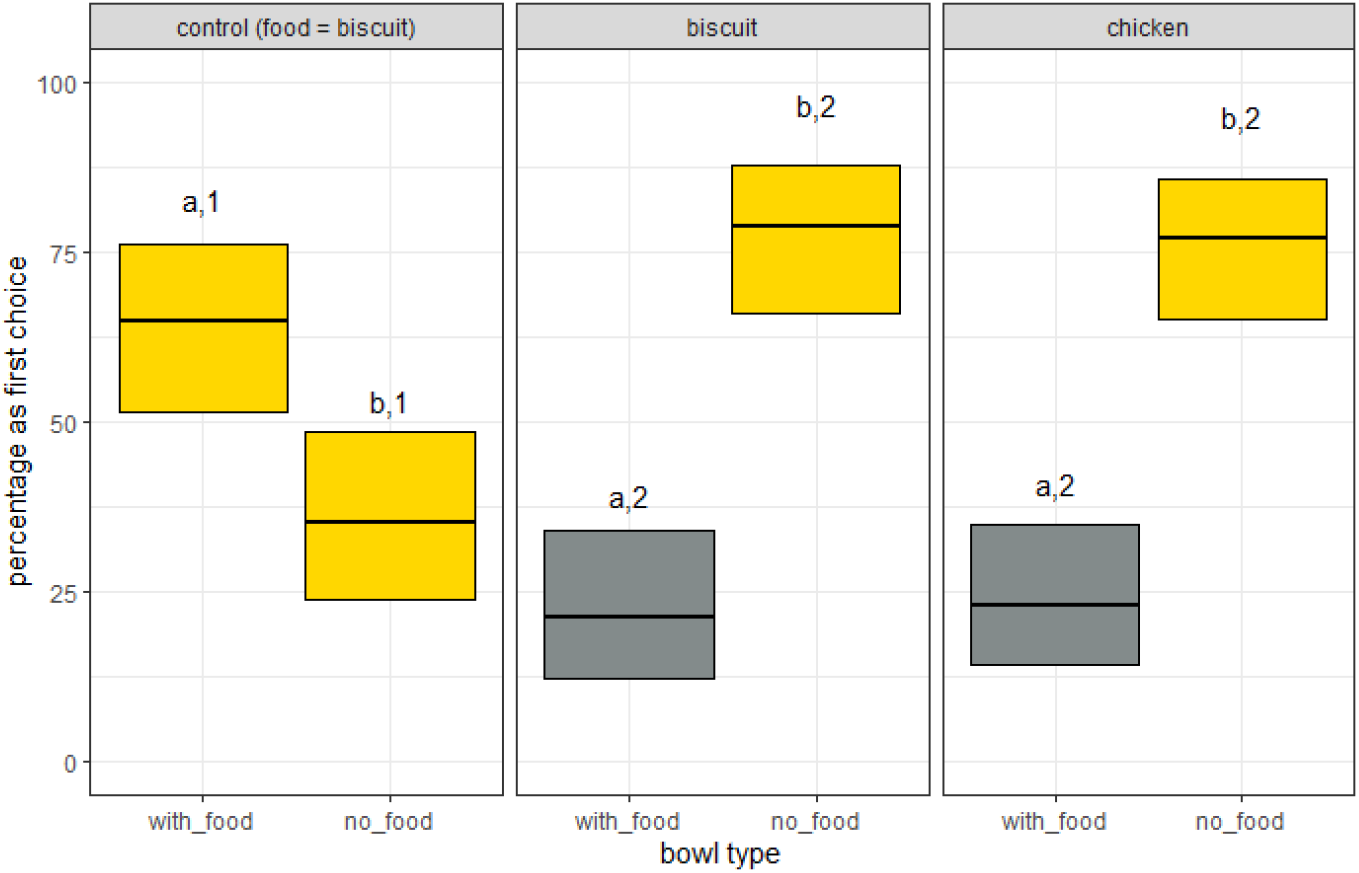
Results of the strength of preference experiments The central bold lines in the boxes are the observations, around which the boxes represent Wilson’s 95% CI. Different letters atop boxes denote significant statistical differences within a condition. Atop boxes, different letters denote significant differences within a condition and different numbers denote significant differences across conditions.

There was no significant difference between the biscuit group and the chicken group (contingency *χ*^*2*^ = 4.05*10^-6, df = 1, p = 0.99), and each of these groups is significantly different than the control group (contingency *χ*^*2*^ tests, control and biscuit: *χ*^*2*^ = 18.819, df = 1, p < 0.001, Carmer’s V = 0.421; control and chicken: *χ*^*2*^ = 18.852, df = 1, p < 0.001, Cramer’s V = 0.404). With Bonferroni correction, the alpha for these three tests was at 0.017.

## 4. Discussion

Around the mid-2010s, India saw a craze of “blue bottles” where people were keeping plastic bottles filled with indigo-dyed water around their houses, in the hope of keeping FRDs away (Jaipuriar, 2016). Masih and Bhadra (2019, unpublished) found that FRDs in and around Kolkata, West Bengal, did not show any specific response towards the bottles. However, people’s attempt at modifying dog behavior by use of color acted as a seed towards our exploration of color preference. The three-choice experiments revealed a preference for yellow over blue or gray in the FRDs. The dogs often explored all the bowls presented in the experiment, and in the case of the two-choice set-up of blue versus gray, they still approached the setup. However, they frequently beelined to yellow even when the gray bowl had food in it and the yellow one did not. Together, these results lead us to the conclusion that the FRDs indeed prefer the color yellow, which is a result of attraction towards yellow, and not due to repulsion towards other colors.

In the three-choice experiment, the presence of food in the bowls did not affect the preferences while anecdotally, the dogs appeared more inclined to participate when food was involved with the setup. Thus, in the two-choice (blue, gray) and control for paint smell experiments only “with_food” trials were done.

Banerjee & Bhadra (2019) previously found Indian FRDs to prefer higher quantities of chicken based on olfactory cues. The dogs were not fooled when deceptive visual cues were used to inflate the amount of chicken present. The same preference however was absent when the food was biscuit, thereby indicating a difference in behavior based on preferability of food (Bhadra et al., 2015). Their study did not consider a 1(present):0(absent) condition for different food amounts. Our control experiment for strength of preference (3.4.1) adds to the findings of Banerjee & Bhadra (2019) and shows that for a 1:0 ratio for biscuits, dogs prefer to choose the option with biscuits. We did not perform the same experiment with chicken since 1:0 is a much higher ratio of chicken than the FRDs are known to discriminate between (Banerjee & Bhadra, 2019).

In the strength of preference experiment with biscuits, we saw that the dogs overwhelmingly chose the empty yellow bowl over the gray bowl that had biscuits. Next, to try and balance the scales, we used chicken, a more attractive food (Bhadra et al., 2015) with the gray bowl. The ∼15g pieces of chicken were the largest that the bowls could comfortably hold without spillage. We tried to maximize the amount of chicken given our current setup, hoping to find a preference shift towards food instead of color, from which we may lower the food amount to find an equilibrium between color and food. Surprisingly, even with the apparent substantial amount of chicken, the preference for yellow dominated. In one trial (Online Resource 4), the focal dog pawed at and attempted to chew the yellow bowl for some time, and then proceeded to solicit from experimenter 1, only returning to obtain the chicken from the gray bowl after a few moments of futile solicitation.

### Possible reasons for the color preference

We do not yet know what exactly is causing this strong preference for yellow. Species confidence (Burley, 1986) might be a possible reason, as many FRDs are shades of orange and brown that will appear yellowish in dog vision (colors simulated by authors using methods from Pongrácz et al., 2017). But while some dogs might appear gray, surely no dog is blue. Since we found no preference between blue and gray, species confidence does not explain the color preference of FRDs. The ecological valence theory (Palmer & Schloss, 2010) links color preference with a color’s usual ecological role or value. Research on innate preference of honeybees (Giurfa et al., 1995) and bumblebees (Raine & Chittka, 2007) had earlier found color preference to be highly correlated with flower colors and associated nectar rewards. For Indian FRDs, most of the food available to them is of human origin (Vanak & Gompper, 2009). Indians often use turmeric (yellow) and dried chili (red) in their food (personal observation), both of which will appear yellow to dogs (Pongrácz et al., 2017). Even raw meat (pink) and blood (red) will appear yellowish (Pongrácz et al., 2017), and if this argument holds, this preference for yellow might predate domestication and be reflected in modern wolves who are the closest relatives to dogs (Bergström et al., 2022). It is important, however, to remember that while scavenging, a dog that is looking for yellow food will come across many false positives: yellow, red, and green human-generated inedible trash. To be accepted, the ecological valence theory must explain how these false positive encounters do not end up diminishing any color preference. It is worth noting that even green grass and trees will appear yellowish to dogs.

Our experiments were limited by the availability of variations in paint color. We finally chose our particular yellow and blue such that their spectral peaks overlapped with respective dog cone cell activity peaks, and thus the hues were of high contrast to the dogs. The brightness of the objects could not be controlled due to our dependence on company-manufactured paints. There are very few and conflicting reports regarding the value of brightness discrimination in dogs (Byosiere, 2018) so we refrain from commenting in that regard. Kasparson et al. (2013) found it easier for pet dogs to discriminate between objects of different hues over objects of the same hue but different brightness. We conjecture brightness to be more important while looking for preference among colors that vary more subtly in hue than in our experiment of high hue contrast.

## 5. Conclusion

Our experiments demonstrate a clear preference for the color yellow over blue and gray in FRDs of India. This preference is so strong that it supersedes the attraction towards food, whether biscuit or chicken. This is the first time that we have observed FRDs ignore a clear food reward in a choice test. Further experiments can help us understand the ecological advantages, if any, of this preference and the reasons behind it. Moreover, comparative studies with pet dogs and wolves can help to understand the evolutionary trajectory of this preference for yellow. The impact of color cues and color preference on training can be explored in the future.

## 6. Acknowledgements

The authors thank Arpan Bhattacharya and Tuhin Shubhra Pal for their participation in the data collection. AR would like to thank Rohan Sarkar and Dr. Udipta Chakraborti for their insights regarding statistical analyses. We thank IISER Kolkata and the Department of Biotechnology, India, for funding support.

## 10. Electronic Supplementary Material

Online Resource 1-5: The videos will be made available when the manuscript is published after peer review.

Online Resource 6:

**Table 1:** First choices for the three-choice experiment across two conditions.

|  | Yellow | Gray | Blue | Total |
| --- | --- | --- | --- | --- |
| with_food | 42 | 18 | 16 | 76 |
| no_food | 30 | 14 | 14 | 58 |
| Total | 72 | 32 | 30 | 134 |

Online Resource 7:

**Table 2:** Distribution of first choice against sexes of the dogs and relative positions of the chosen bowls.

|  | Sex of dogs |  |  | Position of chosen bowl |  |  |  |
| --- | --- | --- | --- | --- | --- | --- | --- |
|  | Male | Female | Total | Left | Center | Right | Total |
| Yellow | 44 | 28 | 72 | 29 | 25 | 18 | 72 |
| gray | 15 | 17 | 32 | 13 | 6 | 13 | 32 |

Online Resource 8:

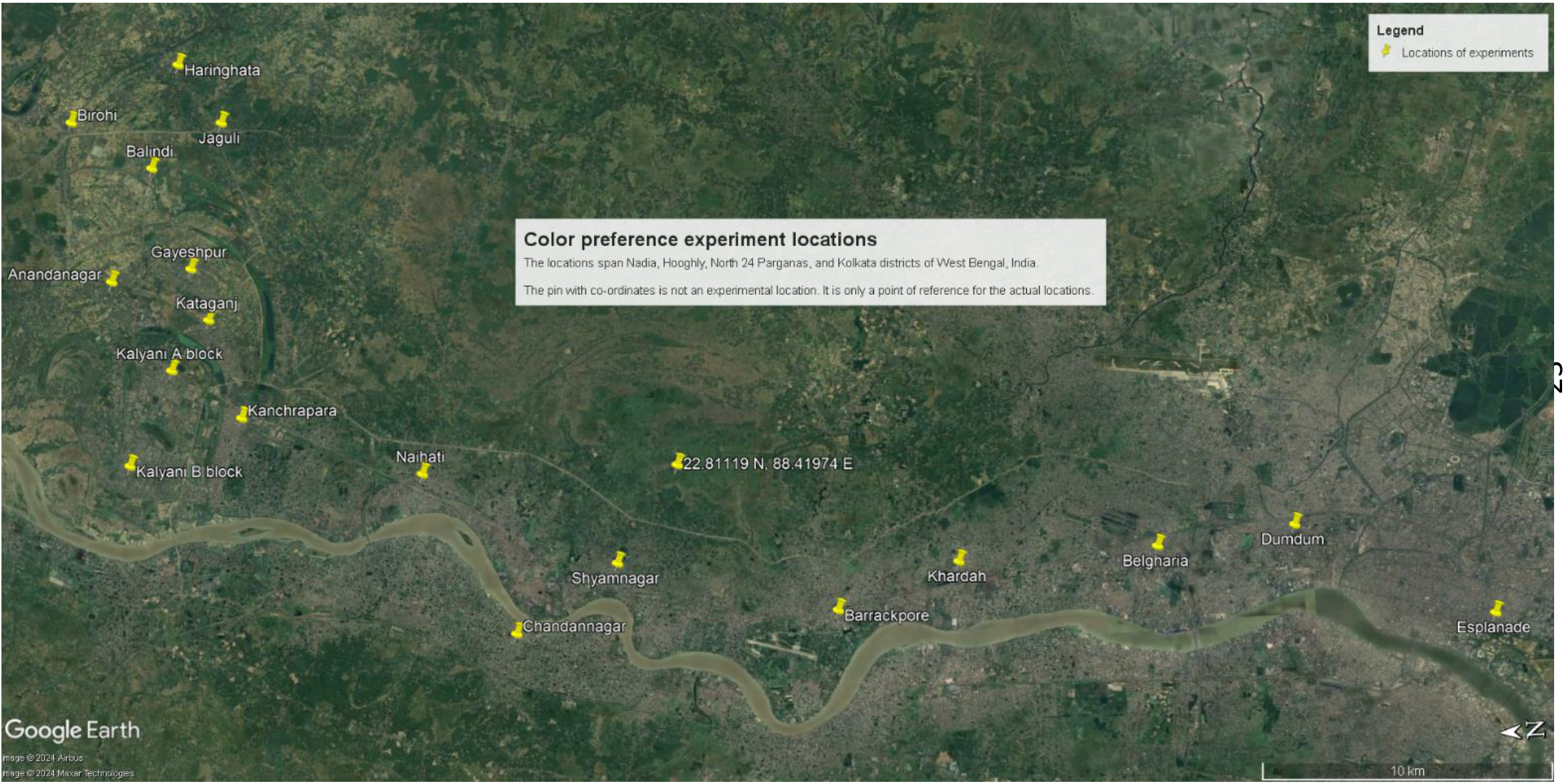

